# Human HMGN1 and HMGN2 are not required for transcription-coupled DNA repair

**DOI:** 10.1101/835868

**Authors:** Katja Apelt, Iris Zoutendijk, Dennis Y. Gout, Diana van den Heuvel, Martijn S. Luijsterburg

**Affiliations:** Department of Human Genetics, Leiden University Medical Center, Einthovenweg 20, 2333 ZC, Leiden, The Netherlands

## Abstract

Transcription-coupled repair (TCR) removes DNA lesions from the transcribed strand of active genes. Stalling of RNA polymerase II (RNAPII) at DNA lesions initiates TCR through the recruitment of the CSB and CSA proteins. The full repertoire of proteins required for human TCR – particularly in a chromatin context - remains to be determined. Studies in mice have revealed that the nucleosome-binding protein HMGN1 is required to enhance the repair of UV-induced lesions in transcribed genes. However, whether HMGN1 is required for human TCR remains unaddressed. Here, we show that knockout or knockdown of HMGN1, either alone or in combination with HMGN2, does not render human cells sensitive to UV light or Illudin S-induced transcription-blocking DNA lesions. Moreover, transcription restart after UV irradiation was not impaired in HMGN-deficient cells. In contrast, TCR-deficient cells were highly sensitive to DNA damage and failed to restart transcription. Furthermore, GFP-tagged HMGN1 was not recruited to sites of UV-induced DNA damage under conditions were GFP-CSB readily accumulated. In line with this, HMGN1 did not associate with the TCR complex, nor did TCR proteins require HMGN1 to associate with DNA damage-stalled RNAPII. Together, our findings suggest that HMGN1 and HMGN2 are not required for human TCR.

## Introduction

Nucleotide excision repair (NER) is a versatile DNA repair system that removes a wide range of helix-distorting DNA lesions from our genome. Many of these DNA lesions, including ultra-violet (UV) light–induced photolesions, as well as lesions inflicted by compounds such as Illudin S and cisplatin potently block transcription^1-3^. Transcription-coupled repair (TCR) is a specialized NER sub-pathway that preferentially removes DNA lesions from actively transcribed DNA strands^4^.

The stalling of elongating RNA polymerase II (RNAPIIo) at DNA lesions was proposed to initiate the TCR pathway by triggering the recruitment of the Cockayne syndrome proteins, CSB and CSA, which, in turn, likely recruit the downstream NER machinery to repair these lesions^5^. Even though the precise molecular mechanism of TCR is far from understood, emerging evidence suggests that modulating chromatin structure is an important prerequisite for mounting an efficient cellular response to transcription-blocking DNA lesions^6-10^.

Packaging of genomic DNA by histone and non-histone proteins into chromatin complicates efficient DNA repair^11^. Even though it could be argued that there is no need for modulating chromatin structure during TCR since chromatin is already rendered permissive to enable transcription^12^, several studies have implicated a role for chromatin-modifying activities associated with TCR-dependent transcription restart in human cells, including histone chaperones FACT and HIRA, and chromatin remodeling factor SNF2H^6-10^. Furthermore, core TCR protein CSB is an ATPase with the ability to remodel nucleosomes *in vitro*^13^, which could contribute to TCR *in vivo*^14^.

Studies in mouse embryonic fibroblasts have revealed that the nucleosome-binding HMGN1 protein has the ability to increase the cellular transcription potential by unfolding higher-order chromatin structure^15^. Interestingly, mouse embryonic fibroblasts deficient in HMGN1 show a decreased repair rate of UV-induced DNA lesions particularly in transcribed genes^16^, suggesting an involvement in modulating chromatin structure during murine TCR. Rather than a specific TCR factor, murine HMGN1 seems to have a more general role in enabling DNA repair in chromatin, since mouse embryonic fibroblasts deficient in HMGN1 show not only defects in the repair of UV-induced lesions^16^, but also in the repair of oxidative DNA lesions^17^, and DNA double-strand breaks^18^.

Most studies addressing the versatile roles of HMGN1 have focused on mouse cells, while our current understanding of the function of HMGN1 in human cells is fairly limited. Although often assumed^5, 12^, experimental evidence showing that HMGN1 has a role in human TCR – similar to its murine counterpart - is lacking. In this study, we established human knockout cells for HMGN1 alone or in combination with HMGN2. Functional analysis revealed that, in contrast to mouse cells, human HMGN1 and HMGN2 are dispensable for human TCR.

## Results

### Generation of human HMGN1 knockout cells

Studies in mouse embryonic fibroblasts have revealed a role of HMGN1 in enhancing the repair rate of UV-induced DNA lesions in particular from transcribed genes ^16^, suggesting a possible involvement of HMGN1 in murine transcription-coupled repair (TCR). However, whether HMGN1 is involved in human TCR has remained unexplored. To study a potential role of HMGN1 in human cells, we used CRISPR/Cas9-mediated genome editing to generate HMGN1 knock-out (KO) cells. To this end, we transfected U2OS cells with vectors encoding HMGN1-specific sgRNAs and Cas9 after which clones were isolated and screened (**Figure 1A**). Western blot analysis using HMGN1-specific antibodies confirmed the knock-out of HMGN1 in two independent clones (**Figure 1A**; clone 2-4 and 2-11). These findings reveal that loss of HMGN1 is viable in human cells and provide a new tool to study the role of HMGN1 in human cells.

**Figure 1.**
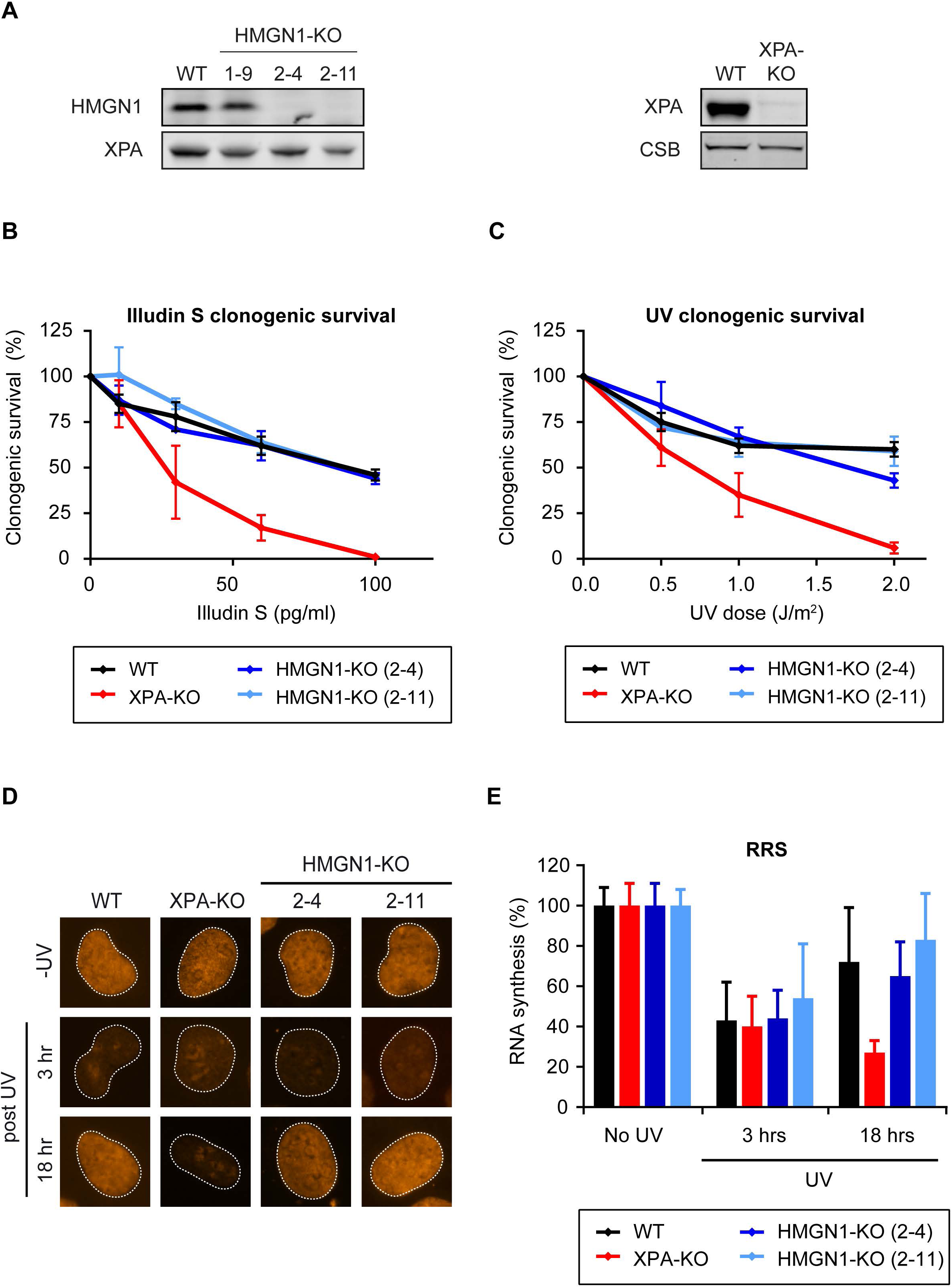
HMGN1 knockout does not impact human TCR. (a) Western blot analysis of U2OS WT and HMGN-KO clones or XPA-KO clone. (b) Clonogenic Illudin S survival or (c) clonogenic UV survival of WT, XPA-KO, and HMGN1-KO cell lines. Data represent mean ± SEM of three independent experiments. (d) Representative microscopy images, and (e) Quantification of RRS after UV on the WT, XPA-KO, and HMGN1-KO cell lines. Data represent mean ± SEM of three independent experiments. Uncropped Western blot data is shown in the Supplementary Information file.

### Human HMGN1-KO cells are resistant to Illudin S and UV

Elongating RNA polymerase II (RNAPIIo) molecules are unable to efficiently bypass DNA lesions that block transcription, including those inflicted by the sesquiterpene drug Illudin S^2^ or by ultra-violet (UV) light, which inflicts photoproducts such as cyclobutane pyrimidine dimers^1^. To overcome the obstacle posed by these DNA lesions, human cells fully depend on TCR to remove these lesions during transcription. Particularly Illudin S-induced lesions are remove exclusively by TCR^2^. To directly compare our HMGN1-KO cells in the same genetic background with TCR-deficient cells, we used CRISPR/Cas9-mediated genome editing to generate XPA knockout (KO) cells. Western blot analysis confirmed the knockout of XPA (**Figure 1A**). Moreover, clonogenic survivals assays in which cells were exposed to increasing concentrations of Illudin S (**Figure 1B**), or increasing doses of UV-C light (**Figure 1C**), confirmed that XPA-KO cells were highly sensitivity to these DNA-damaging agents compared to parental wild-type (WT) cells. In contrast, two independent HMGN1-KO clones, which, were included in parallel, showed no sensitivity to either Illudin S or UV irradiation compared to WT cells (**Figure 1B, C**). These findings show that, in contrast to murine cells^16^, loss of HMGN1 in human cells does not cause sensitivity to transcription-blocking DNA damage.

### Human HMGN1-KO cells show normal transcription recovery after UV

In addition to sensitivity to transcription-blocking DNA damage, another hallmark of TCR-deficient cells is their inability to restart transcription after UV irradiation^19^. To quantify the ability of our HMGN1-KO cells to restart transcription, we performed recovery of RNA synthesis (RRS) experiments. To this end, we either mock treated or exposed cells to UV-C light (6 J/m^2^). Nascent transcripts were pulse-labeled for 1 hour with the cell-permeable thymine analogue 5-ethynyl uridine (5-EU). Nascent transcripts containing 5-EU were visualized via Copper-catalyzed click chemistry of an azide-coupled fluorescent dye. Microscopic analysis revealed that WT cells showed a pronounced inhibition of transcription at 3 hours after UV, due to stalling of RNAPII, while significant transcription restart could be detected at 18 hours after UV irradiation (**Figure 1D, E**). This transcription restart was completely blocked in TCR-deficient XPA-KO cells due to their inability to clear transcription-blocking UV-induced lesions from the genome (**Figure 1D, E**). In contrast, two independent HMGN1-KO clones showed a normal restart of transcription after UV irradiation, suggesting that these cells are not deficient in TCR.

### HMGN2 does not compensate for HMGN1 in TCR

Although HMGN1-deficient mouse embryonic fibroblasts are sensitive to UV irradiation^16^, we did not observe this phenotype in human HMGN1-KO cells. It has been reported that HMGN2 can functionally compensate for HMGN1 in murine cells^20^, and we therefore considered that a similar functional redundancy may mask the role of HMGN1 in human TCR. To test this possibility, we generated HMGN1/HMGN2 double knock-out (dKO) cells by CRISPR-Cas9-mediated genome editing. To this end, U2OS cells were co-transfected with vectors encoding sgRNAs targeting both *HMGN1* and *HMGN2* genes, as well as with a vector encoding the Cas9 protein. Cells were sorted by flow cytometry based on GFP expression encoded on the Cas9 vector, and clones were isolated and screened. Western blot analysis using antibodies specific for human HMGN1 and HMGN2 confirmed the loss of both HMGN proteins in our selected KO clones (**Figure 2A**; clones 1-5 and 1-6). Importantly, two independent HMGN1/HMGN2-dKO clones showed a normal transcription restart after UV irradiation in RRS experiments, while XPA-KO cells, included in parallel failed to resume transcription (**Figure 2B, C**). Furthermore, both HMGN1/HMGN2-dKO clones were resistant to Illudin S-induced DNA lesions, while CSB-KO cells, which were included as a control, were highly sensitive to transcription-blocking lesions induced by this compound (**Figure 2D**). These findings suggest that HMGN2 does not functionally compensate for HMGN1, and that neither HMGN protein is required for TCR in human cells.

**Figure 2.**
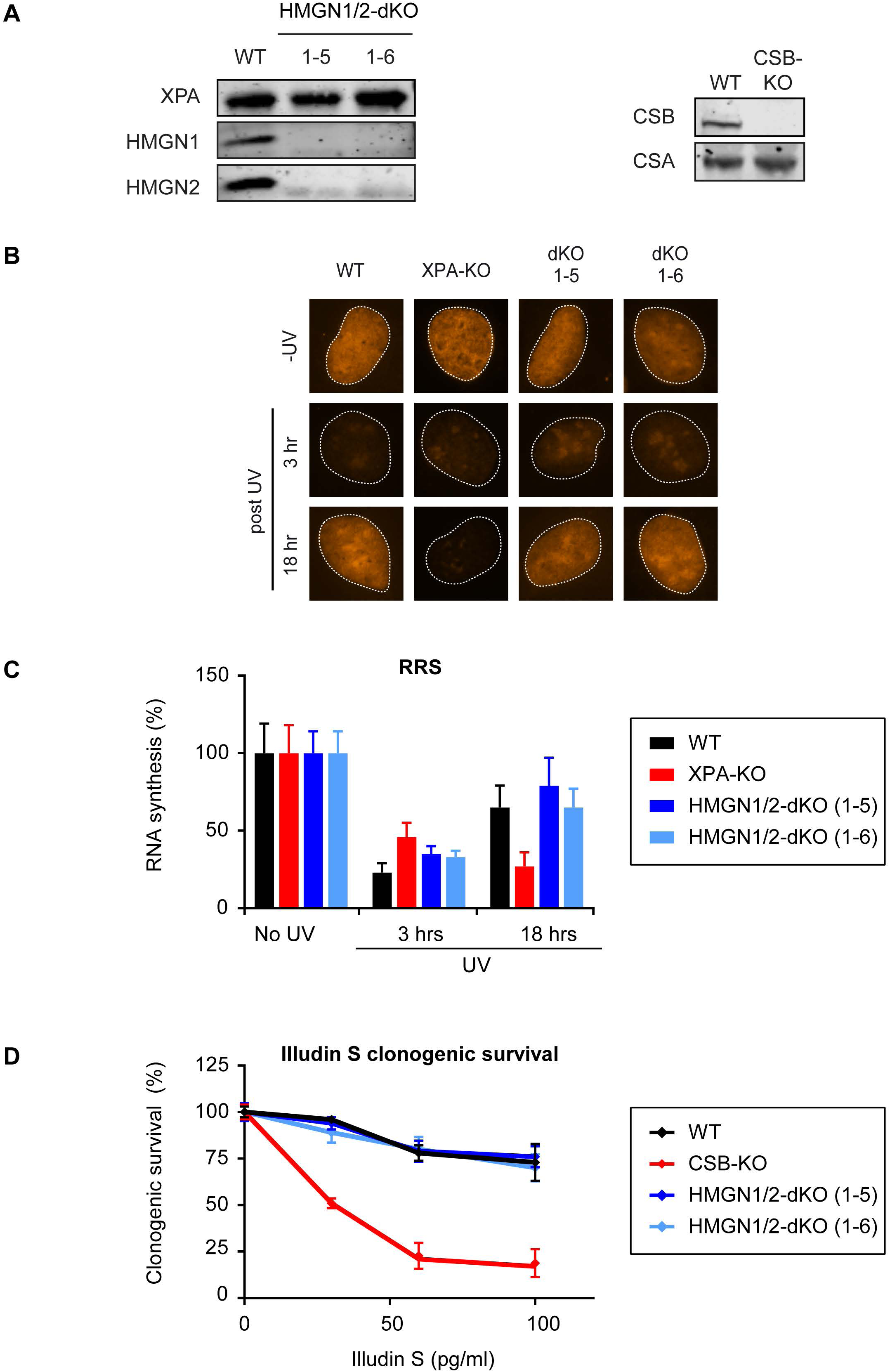
HMGN1 and HMGN2 double knockout does not impact human TCR. (a) Western blot analysis of U2OS WT and HMGN1/HMGN2-dKO clones or CSB-KO clone. (b) Representative microscopy images, and (c) Quantification of RRS after UV on the WT, XPA-KO, and HMGN1/HMGN2-dKO cell lines. Data represent mean ± SEM of five independent experiments. (d) Clonogenic Illudin S survival of WT, CSB-KO, and HMGN1/HMGN2-dKO cell lines. Data represent mean ± SEM of five independent experiments. Uncropped Western blot data is shown in the Supplementary Information file.

### Knockdown of HMGN1 or HMGN2 does not cause TCR defects in human cells

Our previous findings using independently generated HMGN1-KO clones (**Figure 1**), or HMGN1/HMGN2-dKO clones revealed no signs of TCR deficiency (**Figure 2**). To rule out the possibility that these KO cells genetically adapted during their clonal expansion, we decided to employ a more acute way of removing the expression of HMGN proteins. To this end, we employed specific siRNAs to knockdown the expression of the HMGN proteins or XPA as a control. Western blot analysis showed that we achieved efficient knockdown of HMGN1, HMGN2, or XPA within a time-course of four days (**Figure 3A**). However, as observed in our HMGN-KO cells, the knockdown of either HMGN1, HMGN2, or the combined knockdown of both HMGN proteins did not affect the restart of transcription after UV irradiation in RRS experiments, while knockdown of XPA fully impaired this process (**Figure 3B, C**). These findings suggest that acute knockdown of HMGN proteins, like genetic knockout of HMGN proteins, does not cause a deficiency in human TCR.

**Figure 3.**
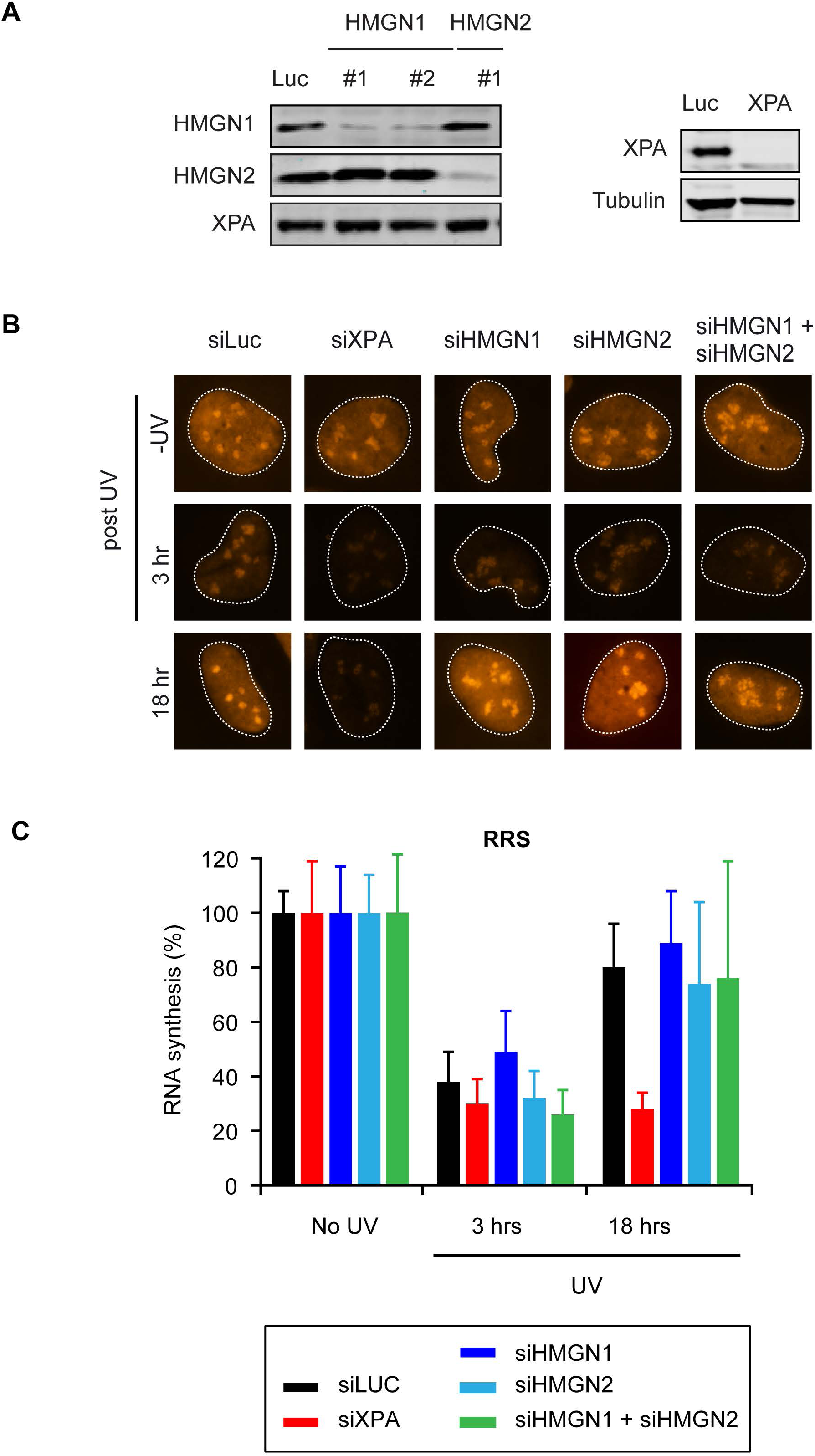
HMGN1 and HMGN2 knockdown does not impact human TCR. (a) Western blot analysis of U2OS cells transfected with the indicated siRNAs. (b) Representative microscopy images, and (c) Quantification of RRS after UV on U2OS cells transfected with the indicated siRNAs. Data represent mean ± SEM of three independent experiments. Uncropped Western blot data is shown in the Supplementary Information file.

### Human HMGN1 does not associate with stalled RNAPII and TCR proteins

Stalling of RNAPII at DNA lesions triggers the association of TCR proteins, including CSB, to initiate repair. We decided to monitor the possible association of HMGN1 with TCR proteins using two independent approaches. We first employed irradiation of cells with a pulsed 266 nm UV-C laser on a live-cell imaging set-up in which all glass optics were replaced by quartz optics to allow full UV-C transmission^21^. To monitor recruitment of proteins using this set-up, we stably expressed either GFP-CSB or HMGN1-GFP in the corresponding KO clones (**Figure 4A**). While UV-C laser-induced DNA damage readily triggered recruitment of GFP-CSB into locally irradiated regions, we failed to detect recruitment of HMGN1-GFP to sites of UV-C laser-induced DNA damage (**Figure 4A**). Importantly, the GFP-tagged human HMGN1 cDNA we used here, was previously shown to complement the phenotype of murine HMGN1-deficient cells demonstrating its functionality ^16^. As an alternative approach, we employed immunoprecipitation under native conditions after UV irradiation using the same cell lines that were used for live-cell imaging. While we could clearly detect a UV-induced association of RNAPIIo after immunoprecipitation of GFP-CSB, we failed to detect the association of endogenous HMGN1 under the same conditions (**Figure 4B**). Similarly, immunoprecipitation of GFP-tagged RNAPII from cells stably expressing GFP-RPB1 ^22^, revealed robust UV-induced interactions with both CSB and CSA (**Figure 4C**), demonstrating that our conditions do allow us to detect interactions with TCR proteins after UV irradiation. However, we could not detect an interaction between GFP-RPB1 and endogenous HMGN1 under these conditions (**Figure 4C**). Reciprocal immunoprecipitation experiments on HMGN1-GFP also did not show any interactions with CSB, CSA, RNAPIIo, or HMGN2 in unirradiated or UV-irradiated cells (**Figure 5A**). Finally, we immunoprecipitated endogenous RNAPIIo, which strongly interacted with CSB, CSA and the TFIIH complex after UV irradiation (**Figure 5B**). However, while all these interactions with RNAPIIo were abolished in CSB-KO cells (**Figure 5B**), in line with the essential role of this protein in TCR, the UV-induced association of these TCR proteins with RNAPIIo was not affected in two independent HMGN1/HMGN2-dKO clones (**Figure 5B**). These findings show that human HMGN1 does not interact with DNA damage-stalled RNAPII and associated TCR proteins, and that both HMGN1 and HMGN2 are dispensable for human TCR.

**Figure 4.**
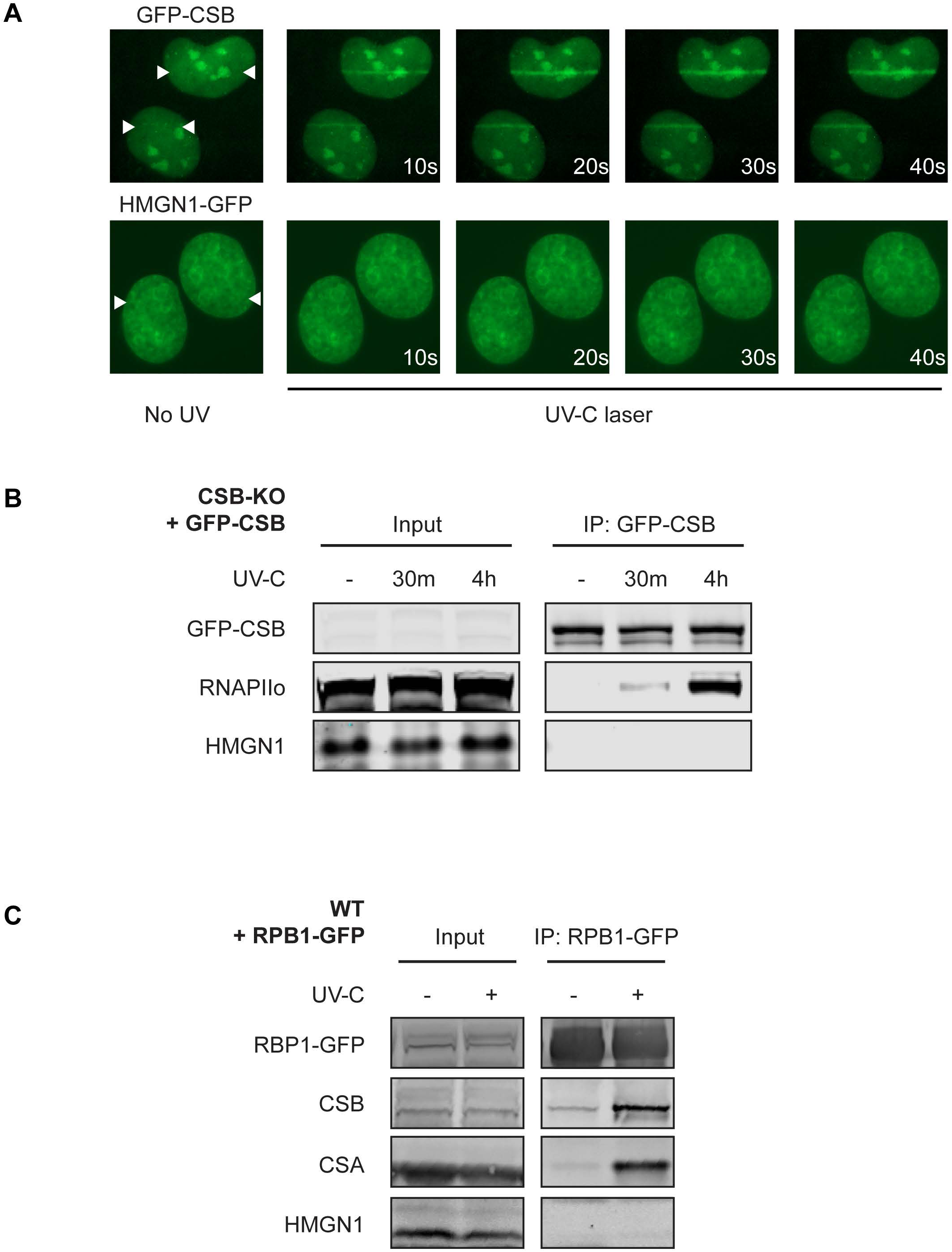
HMGN1 does not associate with the TCR complex. (a) Live-cell imaging on GFP-CSB or HMGN1-GFP after induction of UV-C-laser-induced DNA damage. The position of the laser track is indicated by white arrows. (b) Co-IP of GFP-CSB in unirradiated or UV-irradiated cells at the indicated time-points. (c) Co-IP of RPB1-GFP in unirradiated or UV-irradiated cells. Uncropped Western blot data is shown in the Supplementary Information file.

**Figure 5.**
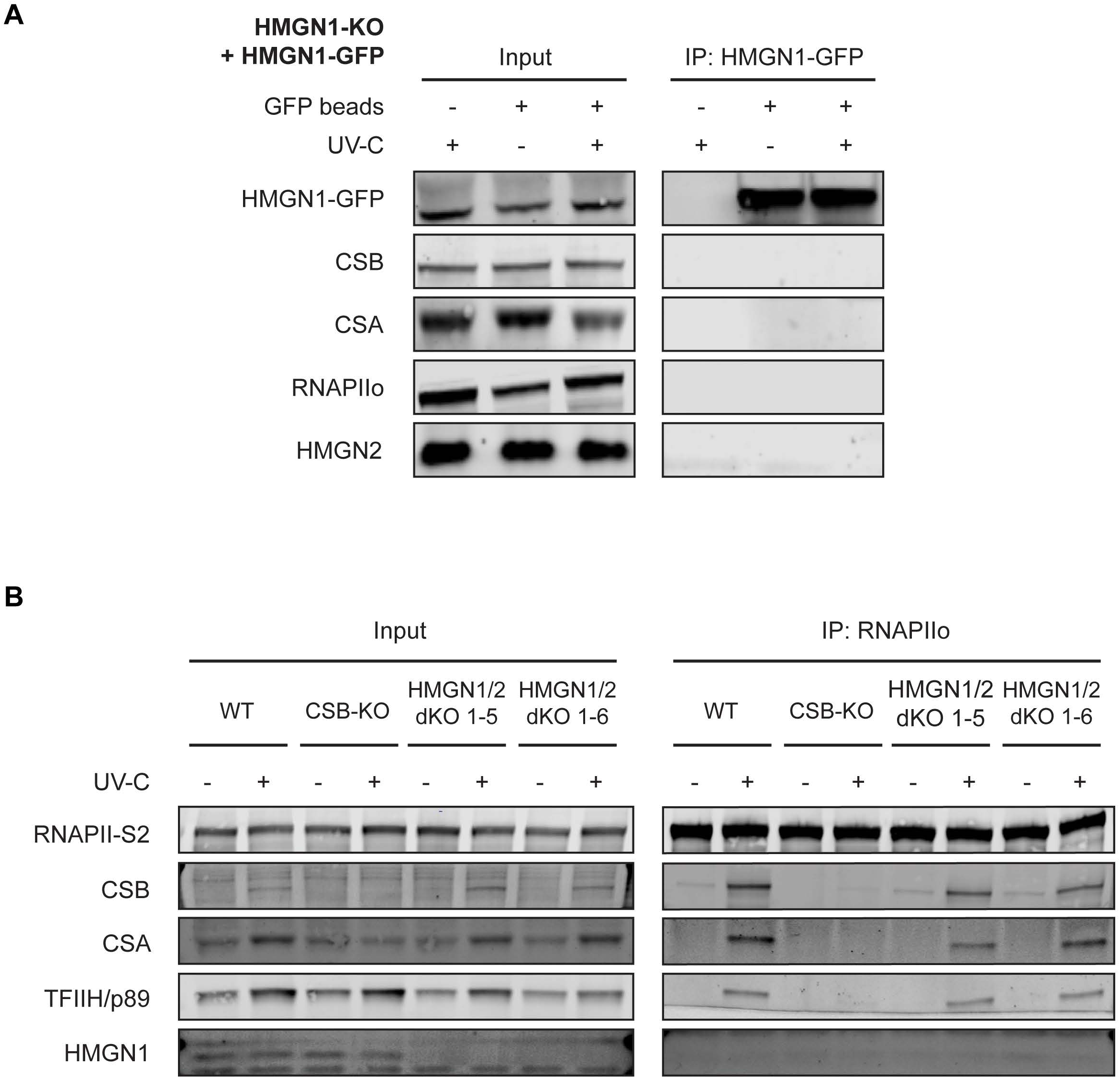
HMGN1 and HMGN2 are not required for TCR complex assembly. (a) Co-IP of HMGN1-GFP in unirradiated or UV-irradiated cells at the indicated time-points. (b) Co-IP of endogenous RNAPIIo in WT, CSB-KO and HMGN1/HMGN2-dKO cell lines in unirradiated or UV-irradiated cells. Uncropped Western blot data is shown in the Supplementary Information file.

## Discussion

Although often inferred, based on studies in mouse embryonic fibroblasts^16^, it is currenly unknown if the nucleosome-binding protein HMGN1 has a role in modulating chromatin structure to enhance transcription-coupled DNA repair (TCR) in human cells. In the current study, we generated human HMGN1 knockout (KO) cells to directly adress this unanswered question. Functional analysis of human HMGN1-KO cells revealed that this nucleosome-binding protein is dispensible for human TCR. Our findings suggest that the role of murine HMGN1 in TCR is not functionally conserved in human cells.

### Human HMGN1 and HMGN2 are not involved in TCR

Several key reviews on TCR mention HMGN1 and list this nucleosome-binding protein as a key factor that modulates human TCR^5, 12, 23^. However, it should be emphasized that while HMGN1-deficient mouse embryonic fibroblasts show decreased repair of UV-induced DNA lesions from active genes^16^, a functional role of HMGN1 in human TCR has not been experimentally adressed. Our results show that knockout or knockdown of the *HMGN1* gene, either alone or in combination with the related *HMGN2* gene, does not impair TCR in human cells. Several functional assays were performed to monitor a functional role in TCR, including clonogenic survival assays after exposure to either Illudin S or UV light, which both trigger transcription-blocking DNA lesions, or recovery of RNA synthesis (RRS) assays, which measure the ability of cells to restart transcription following UV irradiation. While cells knockout for either the *XPA* and *CSB* genes, which are essential for TCR, displayed pronounced defects in all these assays, all the HMGN1-KO or HMGN1/2-dKO clones were indistinguishable from wild-type cells. Thus, our results strongly suggest that HMGN1 and HMGN2 are not required for human TCR.

### Human HMGN1 does not associate with stalled RNAPIIo or TCR proteins

Immunoprecipitation experiments do not support an association of HMGN1 with DNA damage-stalled RNAPIIo, or with either CSB or CSA in response to UV irradiation. These findings are in line with our funtional analysis, and strongly suggest that HMGN1 is not part of the TCR complex. Under the same conditions, we could readily detect a strong UV-induced association between RNAPIIo, CSB, CSA and the TFIIH complex, arguing that our exerimental conditions would allow us to detect the association of HMGN1 if it would occur. Nonetheless, we could neither detect endogenous HMGN1 in CSB precipitates, nor could we detect TCR factors in HMGN1-GFP preciptates. Thus, our interaction experiments do not support an association of HMGN1 with the human TCR complex. Moreover, we show that the association of known TCR factors with DNA damage-stalled RNAPIIo is not affected by the combined loss of the *HMGN1* and *HMGN2* genes.

### Genetic differences between human and murine TCR

Human HMGN1 is a small (100 amino acid) protein with striking sequence conservation (83%) compared to mouse HMGN1 (96 amino acids; **Supplemental Figure 1A**). In fact, the sequence conservation between human and mouse HMGN1 (83%) is far greater than that between human HMGN1 and HMGN2 (47%; **Supplemental Figure 1B, C**). However, our findings suggest that human HMGN1 is not required for human TCR, while a previous study reported that UV-induced lesions in transcribed genes are repaired with decreased efficiency in HMGN1-deficient mouse cells ^16^. Importantly, the UV-sensitive phenotype of these HMGN1-KO mouse cells could be rescued by re-expression of wild-type human HMGN1, but not by mutants that are either unable to bind to nucleosomes, or unable to unfold chromatin, suggesting that this phenotype is a specific effect of the loss of the *HMGN1* gene in mice^16^.

The strong evolutionary similarity (83%) between human and mouse HMGN1 suggest that it is unlikely that the function of HMGN1 between these species is not conserved due to changes at the protein level. However, there are fundamental differences between the organization of TCR in humans compared to mice that are much more likely to underlie these species-specific differences. In humans, the global genome repair (GGR) sub-pathway of NER recognizes and removes UV-induced cyclobutane pyrimidine dimers (CPD) through the DDB2 damage-recognition factor^24-27^. Indeed, inherited mutations in DDB2 cause a predisposition to develop skin cancer^28^. In contrast, mice are largely deficient in the removal of CPDs by GGR owing to very low expression levels of DDB2^29^, and instead rely on TCR to remove CPDs from their genome during transcription. Consequently, TCR-deficient CSB-/- and CSA-/- mice develop skin cancer^30, 31^ due to their inability to repair CPDs, which is not observed in human Cockayne syndrome (CS) patients^32^. Conversely, CS mice do not display strong neurological features^30, 31^, which is a defining hallmark of CS in human patients^32^, further illustrating differences between mice and man when it comes to TCR deficiency. We propose that human cells may not have a need for HMGN1-mediated chromatin modulation to remove CPDs during TCR, because these lesions are targeted by DDB2-mediated GGR. Indeed, several studies have revealed that DDB2 mediates higher-order chromatin unfolding at sites of UV-induced DNA damage^33, 34^ similar to what has been proposed for HMGN1 in mouse cells^15^.

In conclusion, our findings strongly suggest that the role of murine HMGN1 in transcription-coupled DNA repair is not conserved in human cells.

## Experimental Procedures

### Cell lines

All human cells (listed in **table 1**) were cultured at 37°C in an atmosphere of 5% CO2 in DMEM (Thermo Fisher Scientific) supplemented with penicillin/streptomycin (Sigma) and 10% Fetal bovine serum (FBS; Bodinco BV). Parental U2OS (WT) cells were a gift from Andreas Ladurner^35^. U2OS (FRT) cells containing the Flp-In™/T-REx™ system (Thermo Fisher Scientific), were a gift from Daniel Durocher^36^. All cell lines tested negative for mycoplasma contamination.

**Table 1:** Cell lines.

| Cell lines | Origin |
| --- | --- |
| U2OS (FRT) | <sup>36</sup> |
| U2OS (FRT) CSB-KO (1-12) | This study |
| U2OS (FRT) CSB-KO (1-12) + GFP-CSB (3) | This study |
| U2OS (FRT) HMGN1-KO (1-9) | This study |
| U2OS (FRT) HMGN1-KO (2-11) | This study |
| U2OS (FRT) HMGN1-KO (2-4) | This study |
| U2OS (FRT) HMGN1-KO (2-4) + HMGN1-GFP (9) | This study |
| U2OS (FRT) XPA-KO (2-8) | This study |
| U2OS (WT) | <sup>35</sup> |
| U2OS (WT) EGFP-RPB1 | <sup>22</sup> |
| U2OS (WT) HMGN1/HMGN2-dKO (1-5) | This study |
| U2OS (WT) HMGN1/HMGN2-dKO (1-6) | This study |

### Generation of knockout cell lines

To generate single knockouts, U2OS (FRT) cells were co-transfected with pLV-U6g-PPB encoding a guide RNA from the LUMC/Sigma-Aldrich sgRNA library (see **table 2** for plasmids, **table 3** for sgRNA sequences) targeting the *HMGN1, CSB* or *XPA* gene together with an expression vector encoding Cas9-2A-GFP (pX458; Addgene #48138) using lipofectamine 2000 (Invitrogen). Transfected cells were selected on puromycin (1 µg/mL) for 3 days, plated at low density after which individual clones were isolated. To generate HMGN1/HMGN2 double knockouts, U2OS (WT) cells were co-transfected with pLV-U6g-PPB sgHMGN1-2 and pX458 sgHMGN2 also encoding Cas9-2A-GFP. Transfected cells were FACS sorted on BFP/GFP, plated at low density after which individual clones were isolated. Isolated knockout clones were verified by Western blot analysis and/or sanger sequencing. The absence of Cas9 integration / stable expression was confirmed by Western blot analysis.

**Table 2:** Plasmids.

| Plasmids | Origin |
| --- | --- |
| pcDNA5/FRT/TO-Neo | Addgene #41000 |
| pcDNA5/FRT/TO-Puro | This study |
| pcDNA5/FRT/TO-Puro-GFP-N1 | This study |
| pcDNA5/FRT/TO-Puro-HMGN1-GFP | This study |
| pEGFP-N1 | Clontech |
| pHMGN1-EGFP | <sup>16</sup> |
| pLV-U6g-PPB | LUMC/Sigma-Aldrich sgRNA library |
| pLV-U6g-PPB sgCSB | This study |
| pLV-U6g-PPB sgHMGN1-1 | This study |
| pLV-U6g-PPB sgHMGN1-2 | This study |
| pLV-U6g-PPB sgXPA | This study |
| pOG44 | Thermo Fisher |
| pX458 | Addgene #48138 |
| pX458 sgHMGN2-1 | This study |
| pX458 sgHMGN2-2 | This study |

**Table 3:**
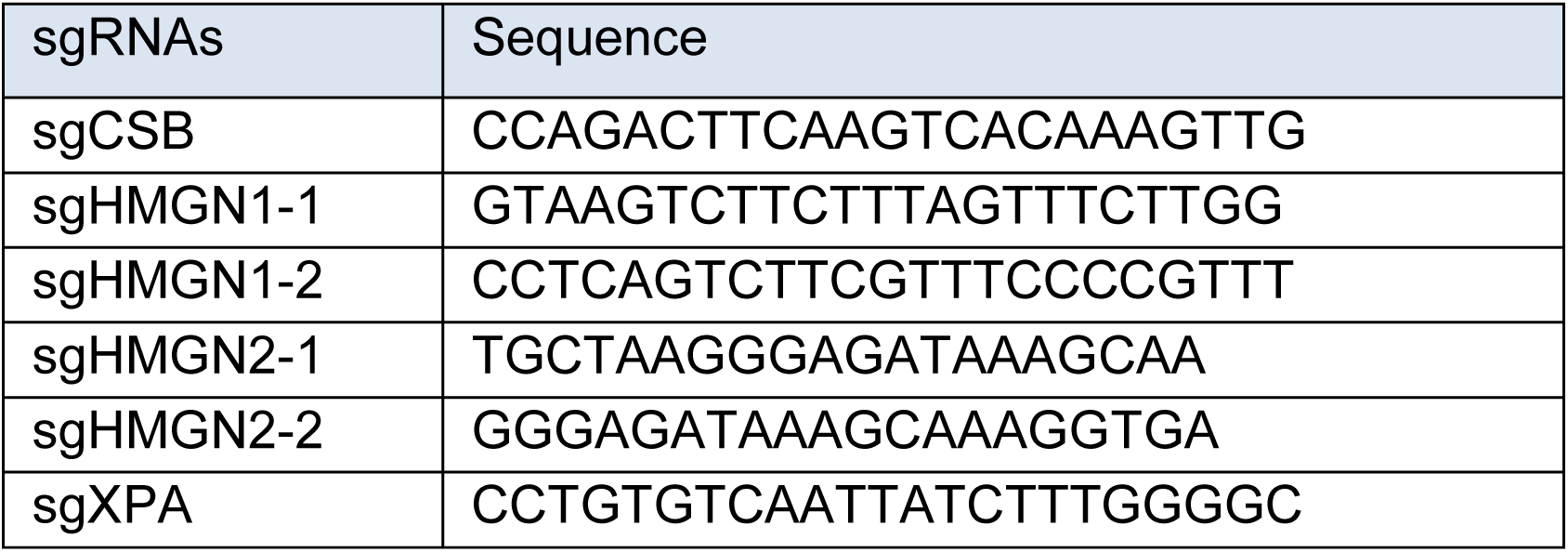
Sequences of sgRNAs.

### Knockdown of HMGN1 or HMGN2

Cells were transfected twice with siRNAs at 0 and 36 hrs and were typically analyzed 60 hrs after the first transfection. All siRNA transfections (see **table 4** for siRNA sequences) were performed with 40 siRNA duplexes using Lipofectamine RNAiMAX (Invitrogen) in OptiMEM without FBS.

**Table 4:**
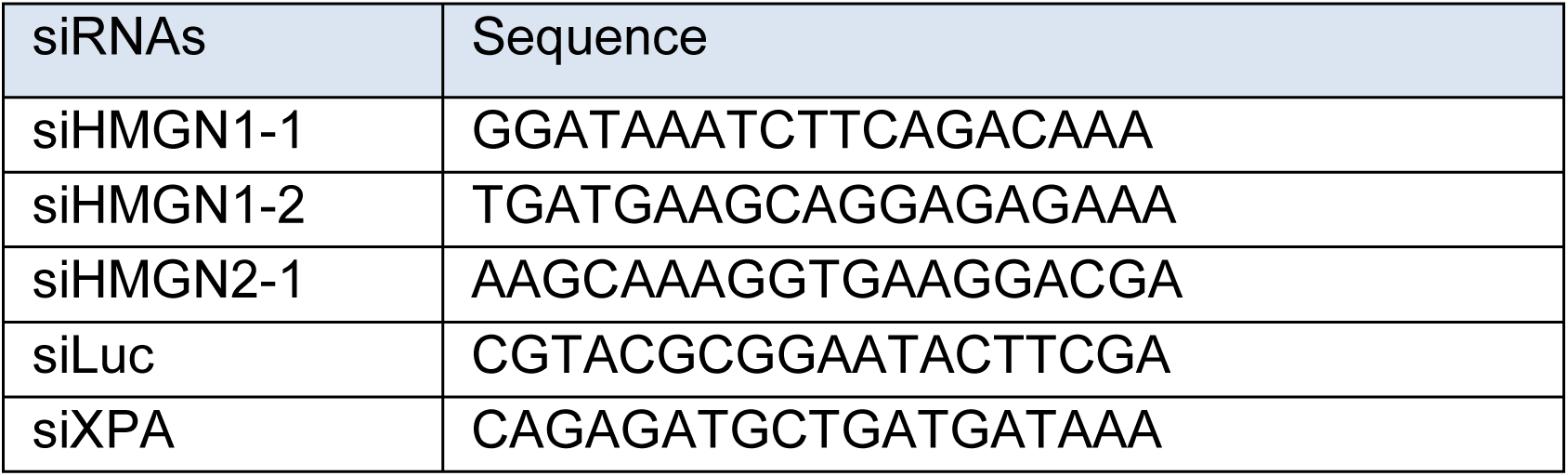
Sequences of siRNAs.

**Table 5:**
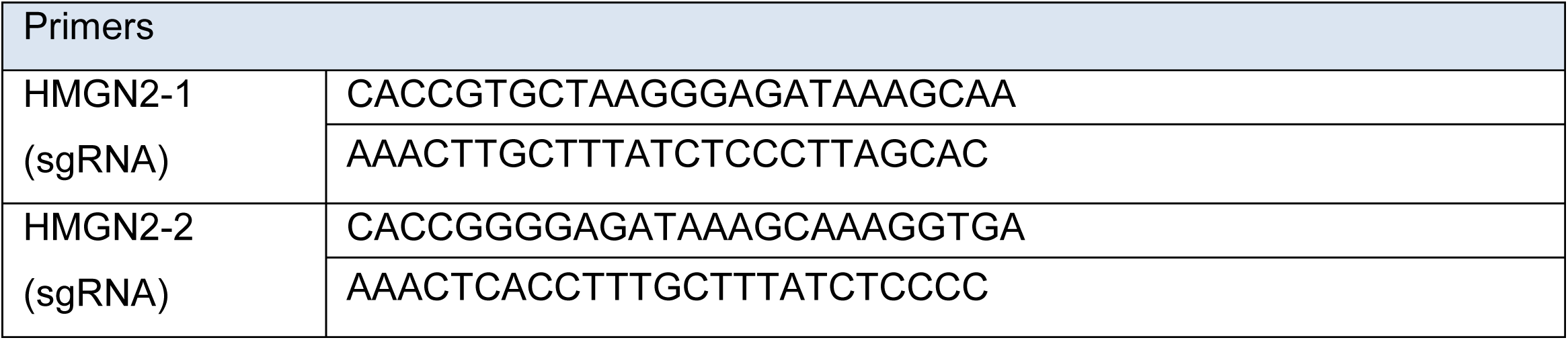
Primers.

### Generation of stable cell lines

Selected knockout clones of CSB and HMGN1 (see **table 1**) were subsequently used to stably express GFP-CSB, or HMGN1-GFP by co-transfection of pCDNA5/FRT/TO-Puro plasmid encoding these genes (5 µg), together with pOG44 plasmid encoding the Flp recombinase (0.5 µg). After selection on 1 µg/mL puromycin and 4 µg/mL blasticidin S, single clones were isolated and expanded. Clones were selected based on their near-endogenous expression level of GFP-tagged proteins compared to parental U2OS Flp-In/T-REx cells. Expression of these GFP-tagged proteins was induced by the addition of 2 µg/ml Doxycycline for 24 hrs.

### Plasmid constructs

To insert sgRNA sequences targeting HMGN2 into pX458, two oligonucleotides (see **tables 3** and **5**) were annealed in annealing buffer (100 mM NaCl, 50 mM HEPES; pH 7,4) by boiling for 5 min in water after which the mixture was allowed to cool down to room temperature. Annealed oligonucleotides were inserted into BbsI-digested pX458 plasmid. The Neomycin resistance gene in pcDNA5/FRT/TO-Neo (Addgene #41000) was replaced with a Puromycin resistance gene. Fragments spanning GFP-N1 (clontech) including the multiple cloning site were inserted into pcDNA5/FRT/TO-puro. The HMGN1 cDNA was inserted as an XhoI / BsrGI fragment into pcDNA5/FRT/TO-Puro-GFP-N1. All sequences were verified by sequencing.

### Clonogenic survival assays

Parental and knockout cell lines were trypsinized, seeded at low density and mock-treated or exposed to an increasing dose of UV light (0.5, 1, 2 J/m^2^ of UV-C 266 nm) or an increasing dose of Illudin S (Santa cruz; sc-391575) for 72 h (15, 30, 60, 100 pg/mL). On day 10, the cells were washed with 0.9% NaCl and stained with methylene blue. Colonies of more than 20 cells were scored. Survival experiments were conducted in triplicate and repeated at least three times.

### Immunoprecipitation for Co-IP

Cells were UV irradiated (20 J/m^2^) or mock treated and harvested 1 h after UV. Chromatin-enriched fractions were prepared by incubating the cells for 20 min on ice in IP-150 buffer (50 mM Tris pH 7.5, 150 mM NaCl, 0.5% NP-40, 2 mM MgCl_2_ with protease inhibitor cocktail (Roche)), followed by centrifugation, and removal of the supernatant. For GFP-IPs, the chromatin-enriched cell pellets were subsequently lysed in IP-150 buffer supplemented with 500 U/mL Benzonase Nuclease (Novagen) for 1 h at 4 °C. For endogenous RNA pol II IPs, the chromatin-enriched cell pellets were lysed in IP-150 buffer supplemented with 500 U/mL Benzonase Nuclease (Novagen) and 2 µg RNAPII-S2 (ab5095, Abcam) for 1 h at 4 °C, followed by adding concentrated NaCl to increase the NaCl concentration to 300mM and incubation of another 30 minutes at 4 °C. Protein complexes were pulled down by 1.5 h incubation with Protein A agarose beads (Millipore; endogenous RNA pol II IPs) or GFP-Trap®_A beads (Chromotek; GFP IPs). For subsequent analysis by Western blotting, samples were prepared by boiling in Laemmli-SDS sample buffer.

### Western blot

Cells were spun down, washed with PBS, and boiled for 10 minutes in Laemmli buffer (40 mM Tris pH 6.8, 3.35% SDS, 16.5% glycerol, 0.0005% Bromophenol Blue and 0.05M DTT). Proteins were separated on 4-12% Criterion XT Bis-Tris gels (Bio-Rad, #3450124) in NuPAGE MOPS running buffer (NP0001-02 Thermo Fisher Scientific), and blotted onto PVDF membranes (IPFL00010, EMD Millipore). Membranes were blocked with blocking buffer (Rockland, MB-070-003) for 2 h at RT, and probed with the indicated antibodies (listed in **table 6**). An Odyssey CLx system (LI-COR Biosciences) was used for detection.

**Table 6:**
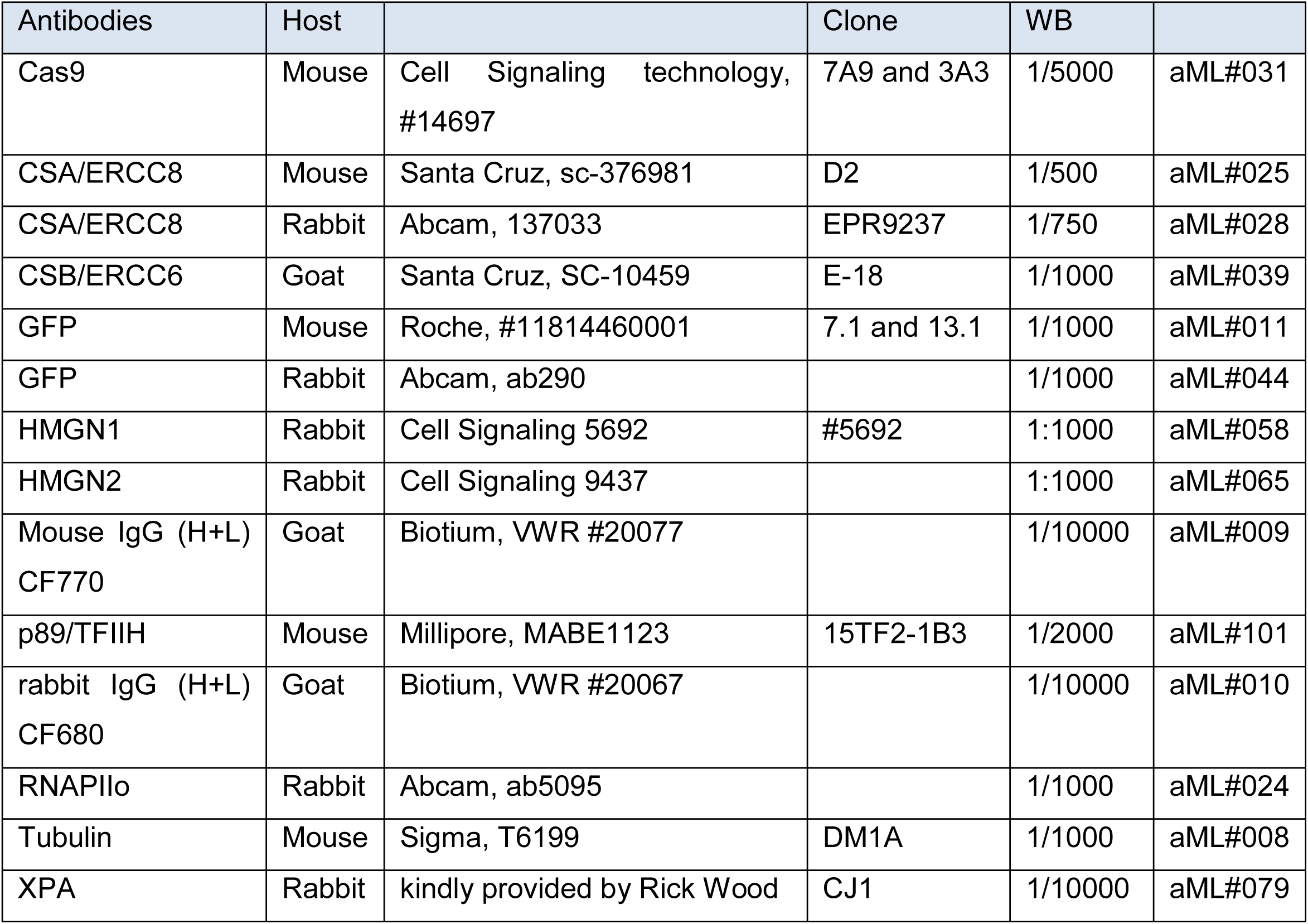
Antibodies.

### RNA recovery assay

30,000 cells were seeded on 12 mm glass coverslips in 24-wells plates in DMEM with 1% FBS. After 24 hours, cells were irradiated with UV-C at a dose of 6 J/m^2^ and incubated in conditioned medium for different time periods (0, 3 and 20 hours) to allow DNA repair and to restart RNA synthesis. Following incubation, nascent RNA was labelled by incubating the cells with 400 μM 5-ethynyluridine (5-EU; Jena Bioscience; CLK-N002-10,), which was then visualized with a click-iT mix consisting of 50mM Tris buffer pH8, 60μM Atto Azide (ATTO-TEC; 647N-101), 4mM CuSO4•5H2O, 10mM L-ascorbic acid (Sigma-Aldrich; A0278) and 1:1000 DAPI (ThermoFisher; D1306) for one hour. Cell were washed three times for 5 minutes with PBS, and mounted on microscope slides (Thermo Scientific) using Aqua Polymount (Polysciences, Inc. #18606). All RRS experiments were conducted in triplicate and repeated at least three times.

### Microscopic analysis of fixed cells

Images of fixed samples were acquired on a Zeiss AxioImager M2 or D2 widefield fluorescence microscope equipped with 63x PLAN APO (1.4 NA) oil-immersion objectives (Zeiss) and an HXP 120 metal-halide lamp used for excitation. Fluorescent probes were detected using the following filters for DAPI (excitation filter: 350/50 nm, dichroic mirror: 400 nm, emission filter: 460/50 nm) and Alexa 555 (excitation filter: 545/25 nm, dichroic mirror: 565 nm, emission filter: 605/70 nm). Images were recorded using ZEN 2012 software and analyzed in Image J.

### UV-C laser microscopy

Cells were grown on 18 mm Quartz coverslips and placed in a Chamlide CMB magnetic chamber in which growth medium was replaced by CO_2_-independent Leibovitz’s L15 medium. Laser tracks were made by a diode pumped solid state 266 nm Yttrium Aluminum Garnet laser (Average power 5 mW, repetition rate up to 10 kHz, pulse length 1 ns) in a UGA-42-Caliburn/2L Spot Illumination system (Rapp OptoElectronic) with laser power set to 20%. This was combined with live cell imaging in an environmental chamber set to 37°C on an all-quartz widefield fluorescence Zeiss Axio Observer 7 microscope, using a 100× 1.2 NA glycerol objective. The laser system is coupled to the microscope via a triggerbox and a neutral density (ND-1) filter is installed to block 90% of the laser light. A HXP 120 V metal-halide lamp was used for excitation.

## Author Contributions

KA generated knockout cells, performed clonogenic survivals, Western blot analysis to validate knockouts, Co-IP experiments, and wrote the paper. IZ generated single knockout cells and double knockout cells, performed clonogenic survivals, Western blot analysis to validate knockouts, and RRS experiments. DG generated double knockout cells, performed clonogenic survivals, Western blot analysis to validate knockouts, RRS experiments, and UV-C laser experiments. DvdH generated knockout cells, and Co-IP experiments. MSL performed UV-C laser experiments, supervised the project, and wrote the paper.

## Acknowledgments

The authors acknowledge Rick Wood for his generous gift of antibodies. Andreas Ladurner provided U2OS (WT) cells, and Dan Durocher provided U2OS (FRT) cells. This work was funded by an LUMC Research Fellowship and an NWO-VIDI grant (ALW.016.161.320) to MSL.

## Competing interests

The authors declare no conflict of interest.

## Data availability

The datasets generated during and/or analysed during the current study are available from the corresponding author on reasonable request.

---

**Supplemental Figure 1.**
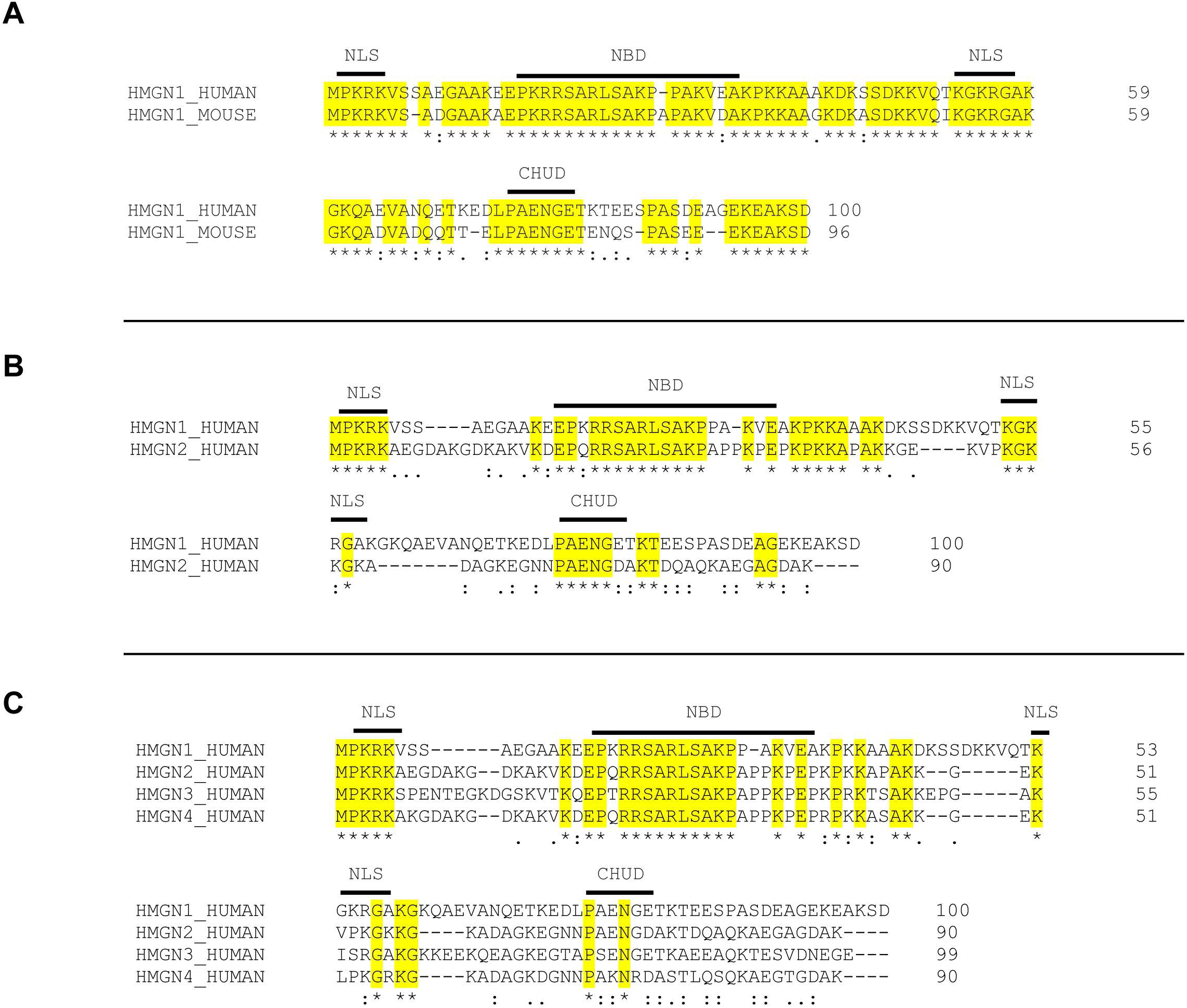
Alignment of HMGN proteins. (a) Sequence alignment of human and mouse HMGN1 proteins. (b) Sequence alignment of human HMGN1 and HMGN2 proteins. (C) Sequence alignment of human HMGN1, HMGN2, HMGN3, and HMGN4 proteins. All sequences were aligned with ClustalW. NLS = nuclear localization signal, NBD = nucleosome-binding domain, CHUD = chromatin-unfolding domain.

