## Supplemental File 1 for "Human HMGN1 and HMGN2 are not required for transcription-coupled DNA repair"

**Supplementary Information file**

Figure 1A

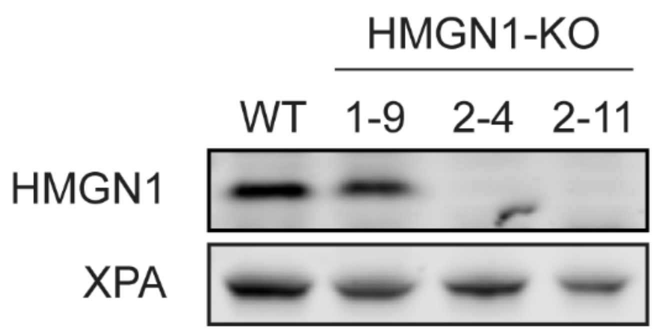


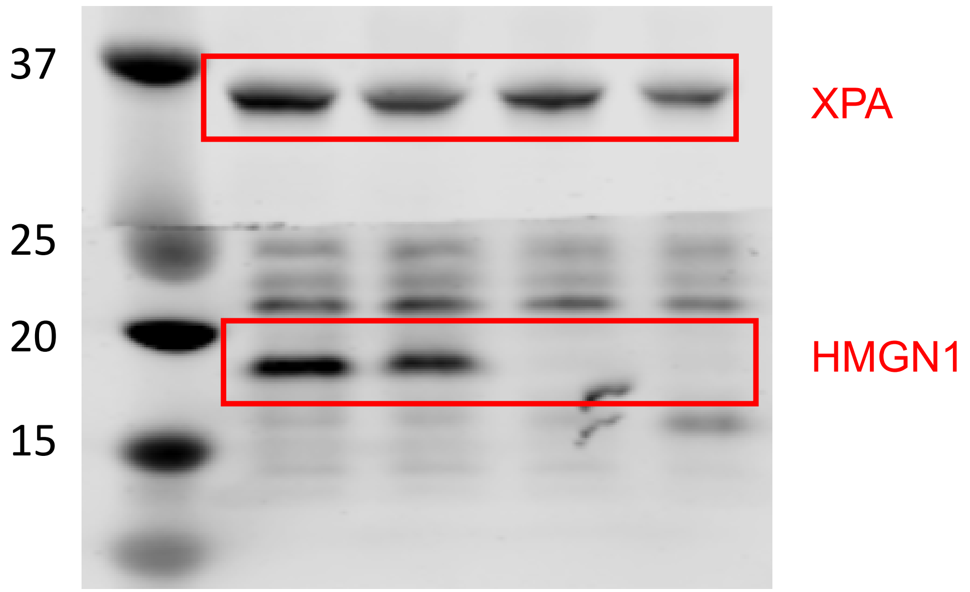

Figure 1A

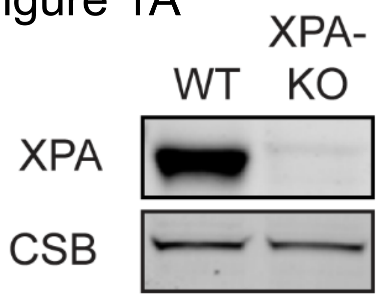


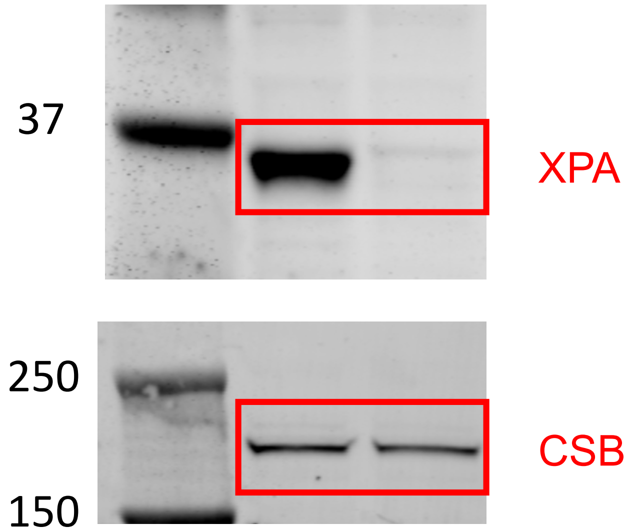

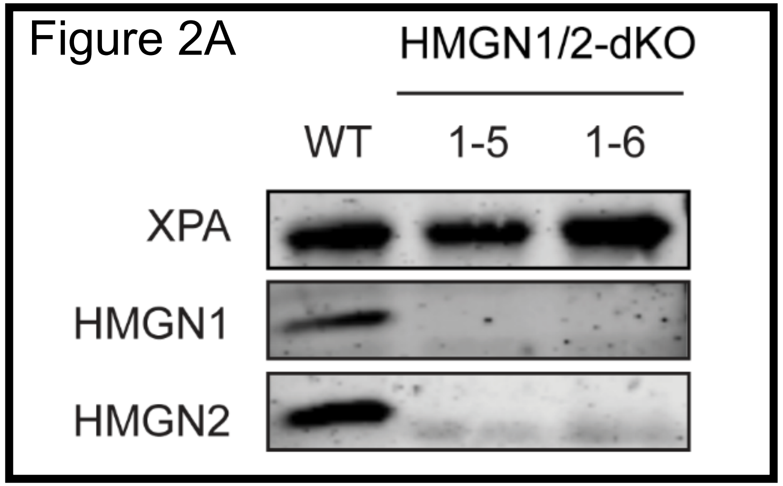
Figure 2A
HMGN1/2-dKO
WT
1-5
1-6
XPA
HMGN1
HMGN2

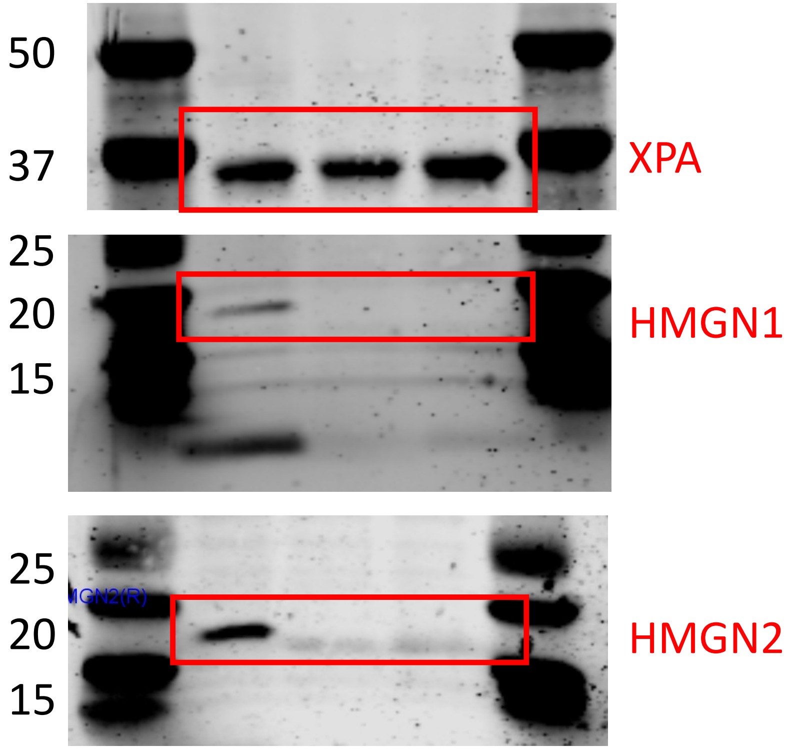
50
37
XPA
25
20
15
HMGN1
25
20
15
HMGN2

Figure 2A

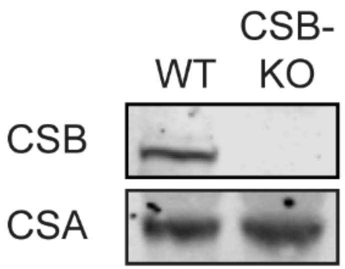


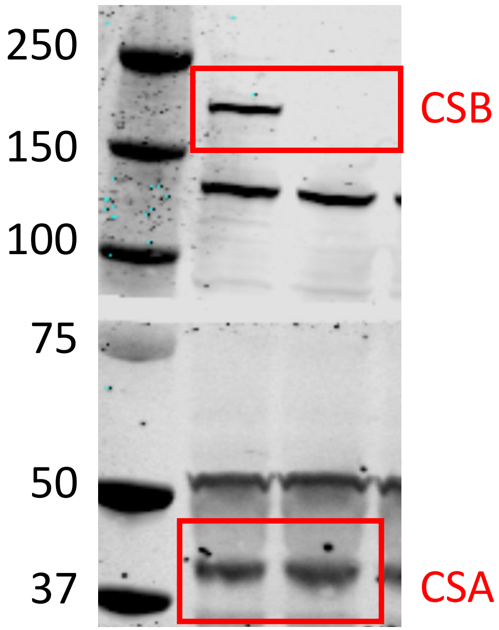

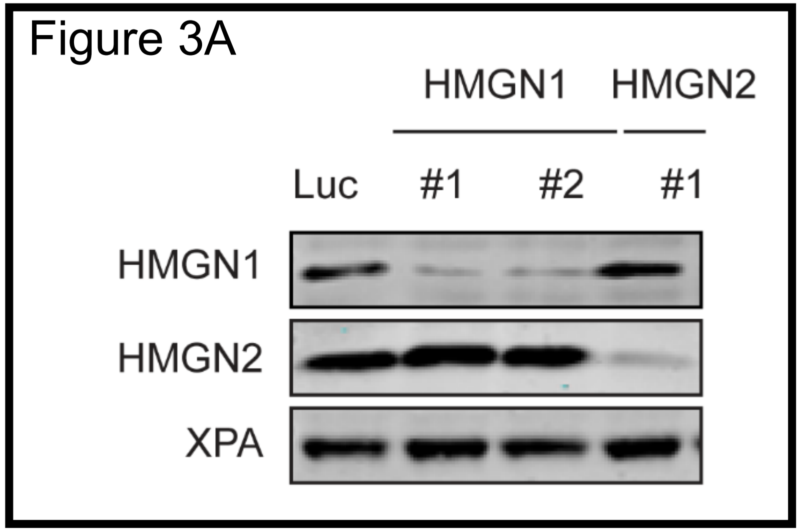
Figure 3A
HMGN1
HMGN2
Luc
#1
#2
#1
HMGN1
HMGN2
XPA

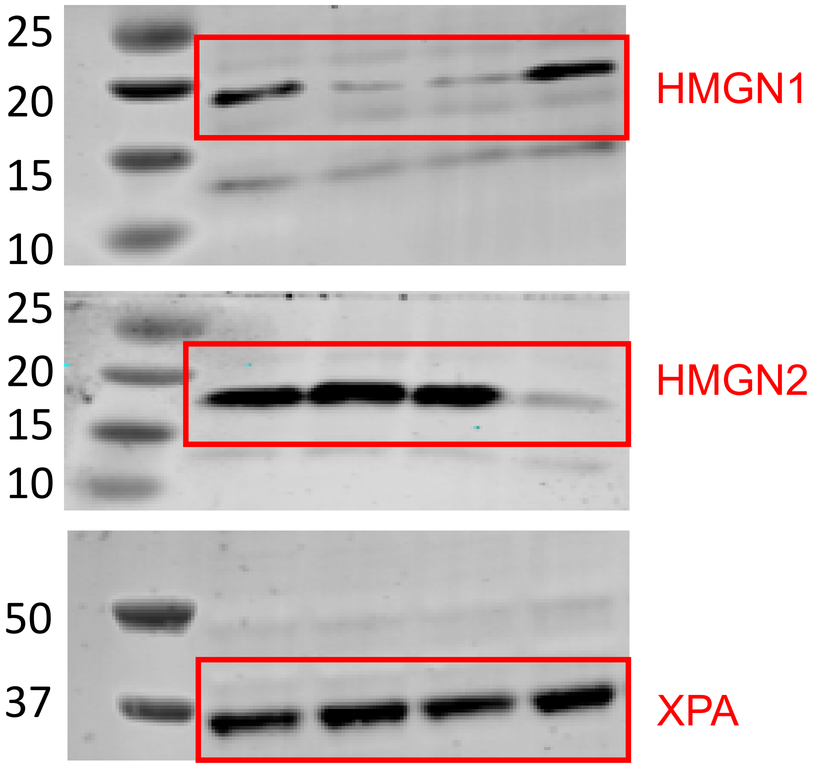
25
20
15
10
HMGN1
25
20
15
10
HMGN2
50
37
XPA

Figure 3A

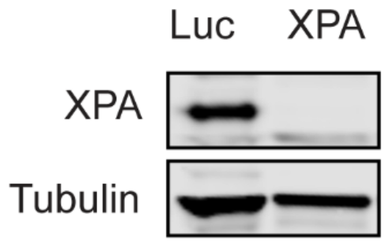


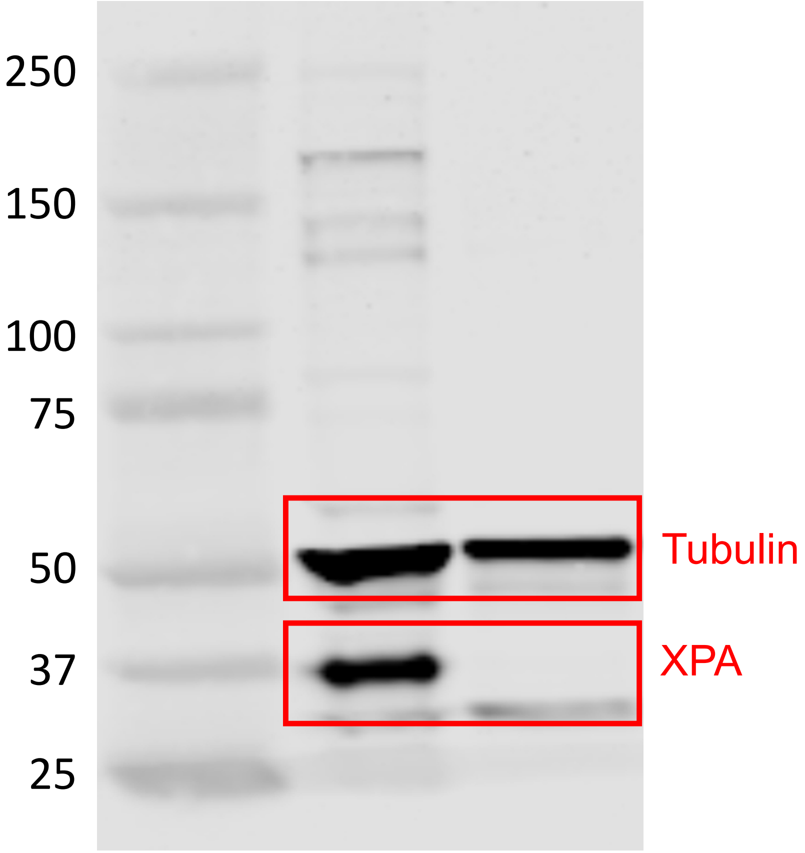

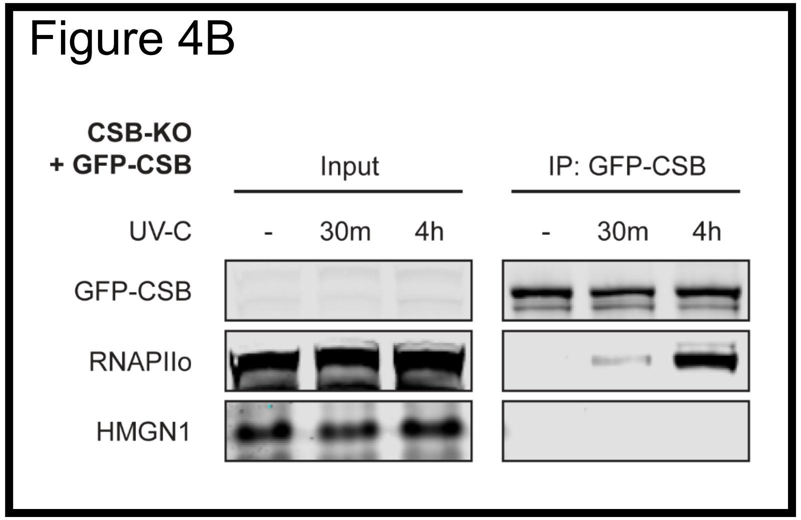
Figure 4B
CSB-KO
+ GFP-CSB
Input
IP: GFP-CSB
UV-C
-
30m
4h
-
30m
4h
GFP-CSB
RNAPIIo
HMGN1

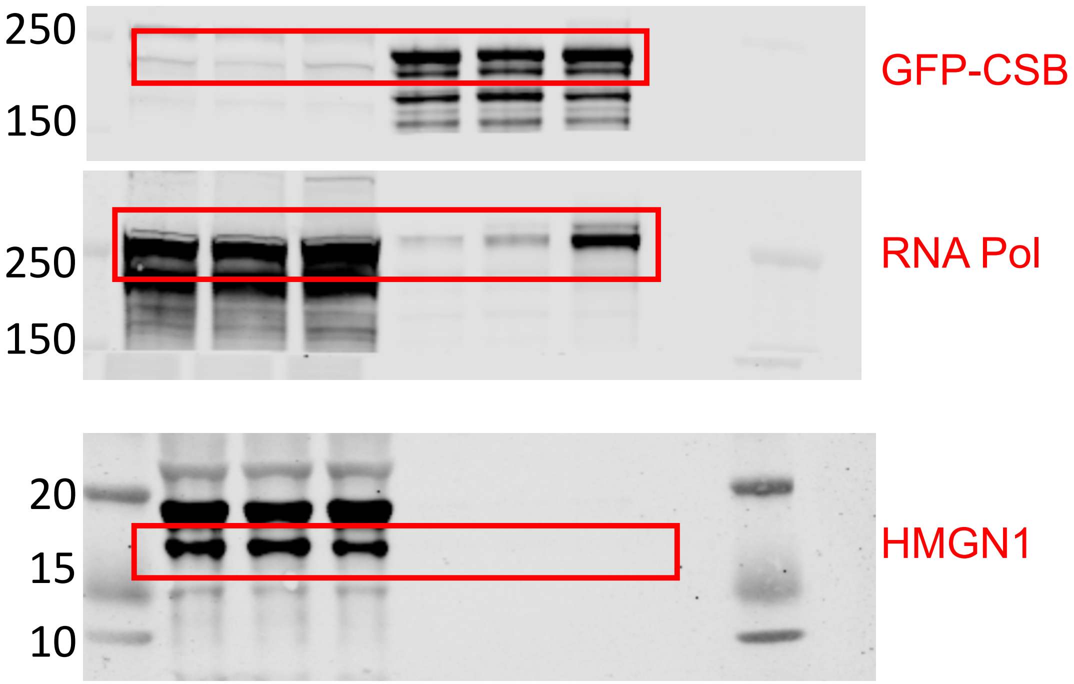
250
150
GFP-CSB
250
150
RNA Pol
20
15
10
HMGN1

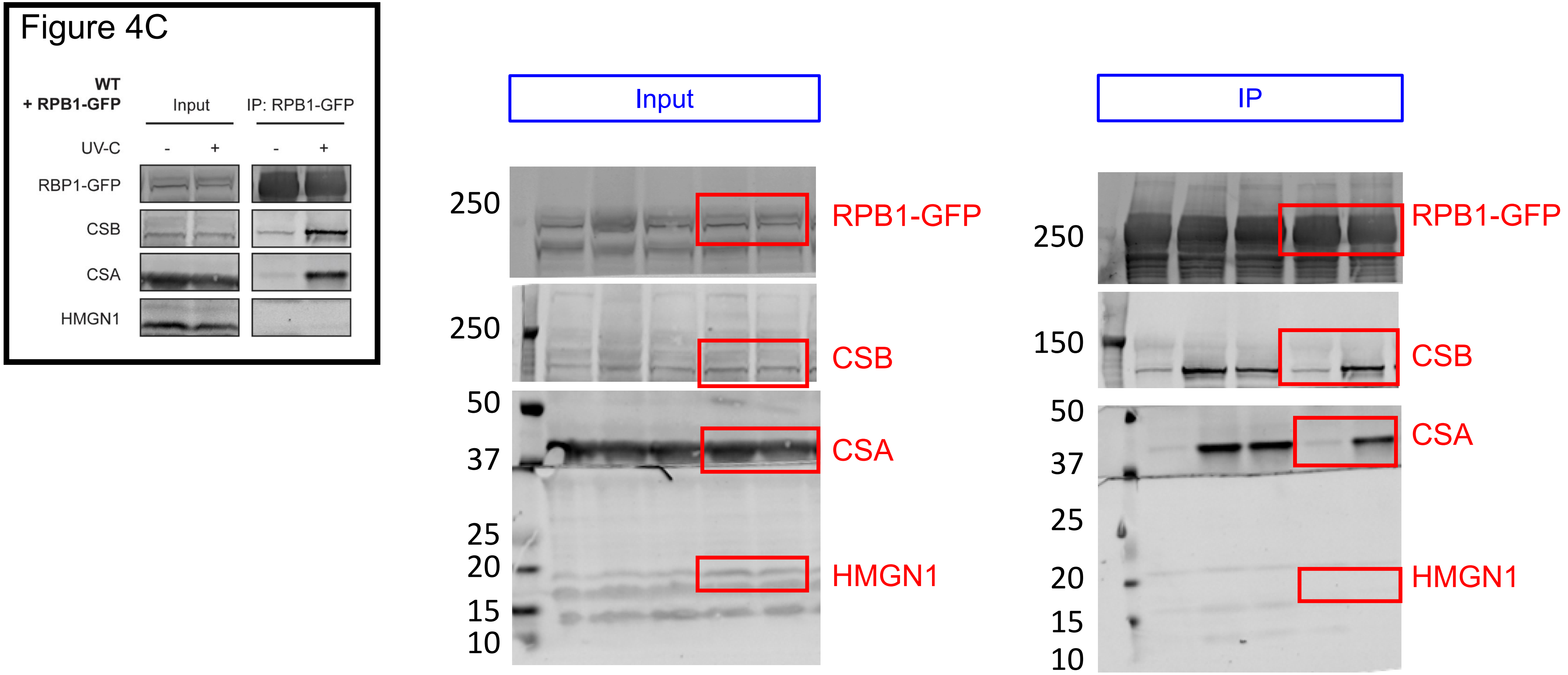
Figure 4C
WT
+ RPB1-GFP
Input
IP: RPB1-GFP
UV-C
-
+
-
+
RBP1-GFP
CSB
CSA
HMGN1
Input
IP
250
250
50
37
25
20
15
10
RPB1-GFP
CSB
CSA
HMGN1
250
150
50
37
25
20
15
10
RPB1-GFP
CSB
CSA
HMGN1

### Figure 5A

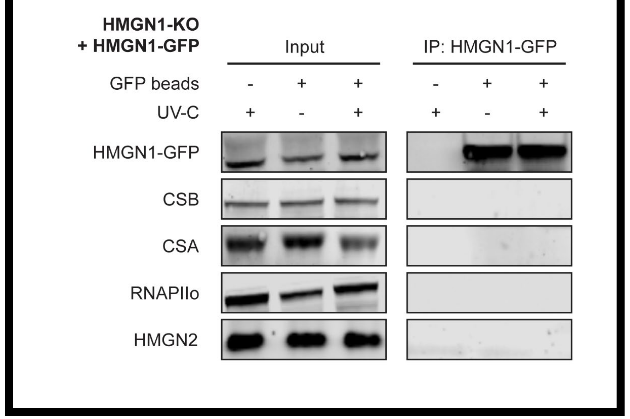
HMGN1-KO
+ HMGN1-GFP
Input
IP: HMGN1-GFP
GFP beads - + + - + +
UV-C + - + + - +
HMGN1-GFP
CSB
CSA
RNAPIIo
HMGN2

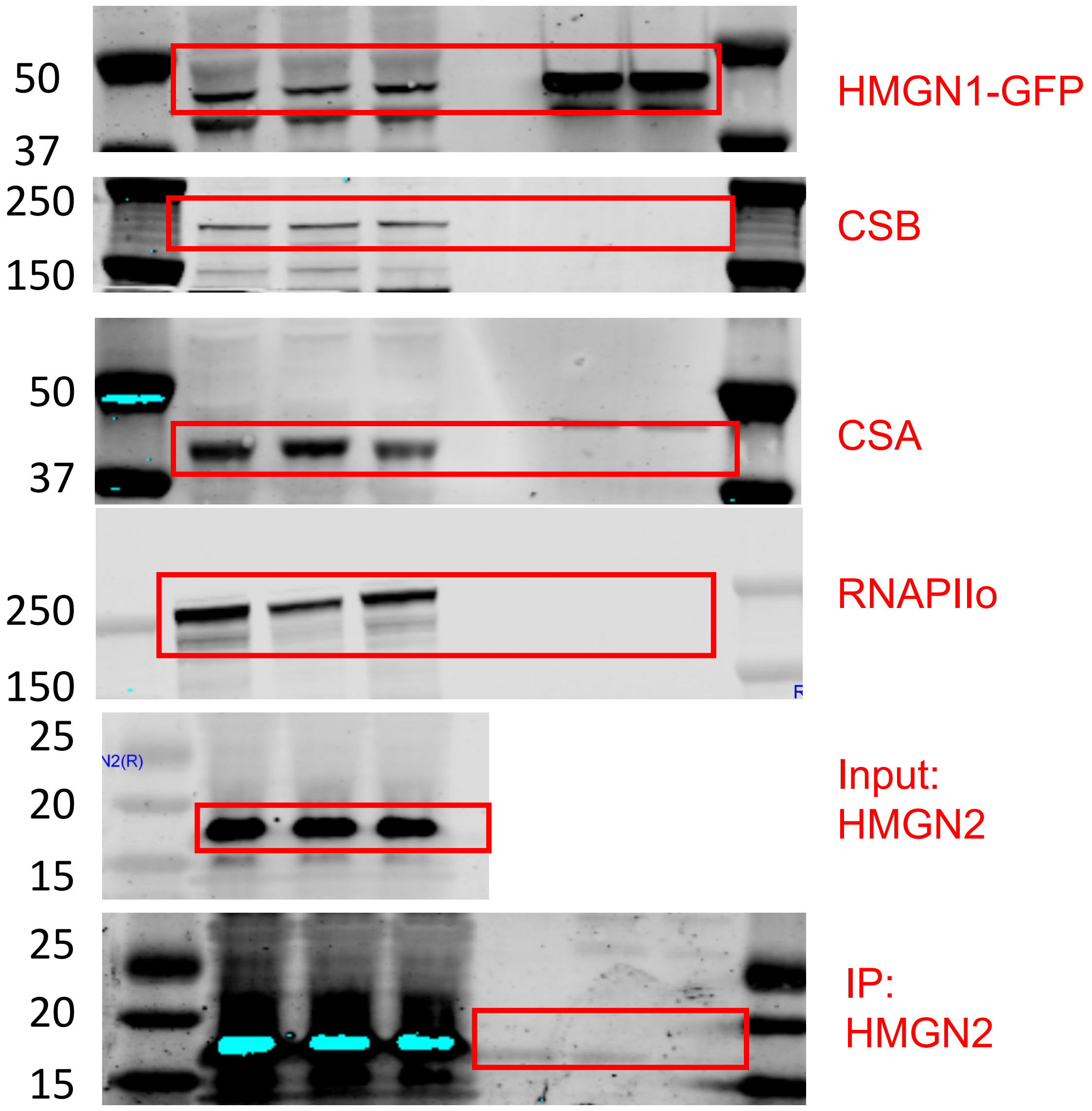
50
37
HMGN1-GFP
250
150
CSB
50
37
CSA
250
150
RNAPIIo
25
20
15
Input:
HMGN2
25
20
15
IP:
HMGN2

Figure 5B

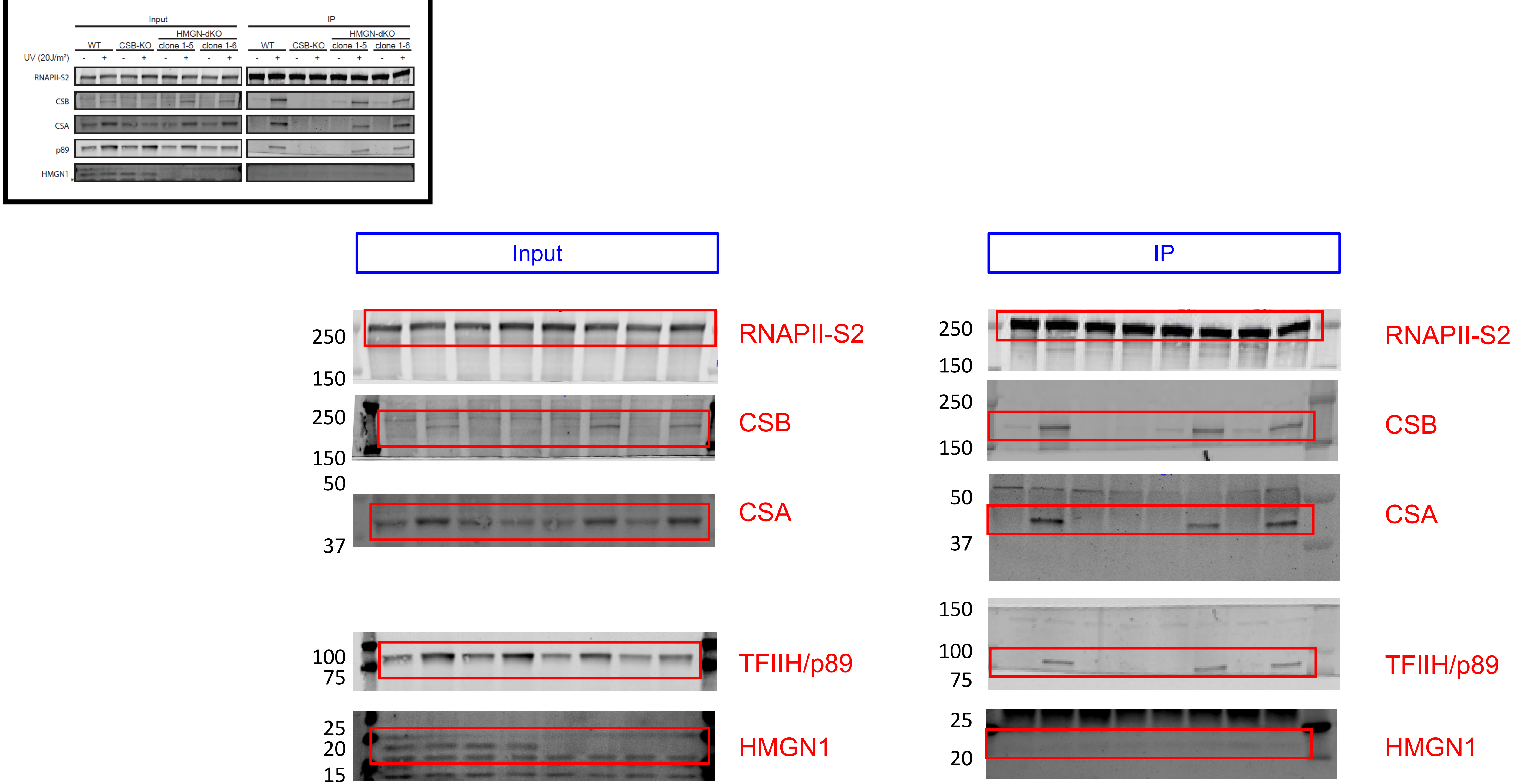
Input
IP
WT
CSB-KO
HMGN-dKO
clone 1-5
clone 1-6
UV (20J/m²)
RNAPII-S2
CSB
CSA
p89
HMGN1
TFIIH/p89
250
150
50
37
100
75
25
20
15
